# Residence time structures microbial communities through niche partitioning

**DOI:** 10.1101/2024.08.22.609267

**Authors:** Emmi A. Mueller, Jay T. Lennon

## Abstract

Much of life on Earth is at the mercy of currents and flow. Residence time (τ) estimates how long organisms and resources stay within a system based on the ratio of volume (*V*) to flow rate (*Q*). Short residence times promote immigration but may prevent the establishment of species that cannot quickly reproduce, or resist being washed out. In contrast, long residence times reduce resource input, selecting for species that can survive on a low supply of energy and nutrients. Theory suggests that these opposing forces shape the abundance, diversity, and function of flowing systems. In this study, we subjected chemostats inoculated with a complex lake microbial community to a residence time gradient spanning seven orders of magnitude. Microbial abundance, richness, and evenness increased with residence time, while functions like productivity and resource consumption decreased along the gradient. Microbial taxa were non-randomly distributed, forming distinct clusters of short-τ and long-τ specialists, reflecting a pattern of niche partitioning. Consistent with theoretical predictions, we demonstrate that residence time shapes assembly processes with direct implications for biodiversity and community function. These insights are crucial for understanding and managing flowing environments, such as animal gut microbiomes, soil litter invertebrate communities, and plankton in freshwater and marine ecosystems.

## INTRODUCTION

In nature, physical forces influence the movement of resources and organisms in complex ways that determine the structure and function of communities (Kadlec 1994, Bell et al. 2002, Ambrosetti et al. 2003, Kashyap et al. 2013). Residence time (τ) captures some of this physical complexity by estimating the average amount of time that passively moving particles and organisms spend in a system. Operationally, it is quantified as the ratio of volume (*V*) to flow rate (*Q*) of a system (Nauman 2008). Residence time has long been used to characterize the growth and metabolism of planktonic organisms in continuous flow reactors (i.e., chemostats).

For example, residence time theory predicts that the size (*N*) and growth rate (*µ*) of a population are both maximized when the dilution rate of a system (1/τ) is equal to the maximum growth rate (*µmax*) of the population (Smith and Waltman 1995). Many of the core principles derived from residence time form the basis of consumer-resource theory in ecology (Tilman 1982; Sommer 1992; Grover 1997).

Despite differences in scale and complexity, common physical processes underlie the population- and community-level patterns that might emerge in systems with varying residence times. Specifically, two opposing forces influence survival and reproduction along a residence time gradient. When τ is short, there should be high rates of immigration from the regional pool of species along with an elevated resource supply that should favor fast-growing species.

Although some individuals are likely to be washed out of the system at high flow rates, slower-growing individuals may resist this source of mortality through attachment to surfaces or active dispersal against currents (Stemmons and Smith 2000). In contrast, when τ is long, resource supply rate may be insufficient to meet the maintenance requirements for some species due to resource limitation. Under such conditions, individuals that do not invest in persistence strategies, such as slow growth, efficient resource consumption, or decreased metabolism, are likely to starve (Lennon and Jones 2011, Shoemaker et al. 2021). Species may contend with the opposing forces of washout and resource limitation in various ways depending on their traits and assembly processes (Locey and Lennon 2019), which could potentially lead to niche partitioning along the residence time axis.

While residence time is a powerful framework that is well-supported by empirical observations (Smith and Waltman 1995, Amster et al. 2020), it was developed for idealized conditions that are commonly violated in nature. For example, residence-time theory typically assumes no immigration, constant growth of a population, and uniformly mixed conditions that support simple communities consisting of just a few species (Harmand et al. 2017). Recently, efforts have been made to incorporate additional complexity into residence time theory using stochastic individual-based models (IBMs). These models investigate community responses to residence time in a one-dimensional environment with a limited regional pool of species and resources (Locey and Lennon 2019). Across combinations of volumes and flow rates, the models identify potential constraints while making predictions that organismal abundance (*N*), species richness (*S*), and productivity should peak at intermediate residence time (Locey and Lennon 2019). Although these models capture multiplicative interactions among species and resources, they are still relatively simple compared to the complexities that can emerge in real world systems.

Here, we experimentally evaluated the effects of residence time on measures of community structure and function in continuous flow reactors (i.e., chemostats) exposed to a residence time gradient spanning seven orders of magnitude. We inoculated these chemostats with an assemblage of lake microorganisms and characterized patterns in abundance and productivity that developed in the chemostats. In addition to assessing community assembly and resource consumption, we tested for patterns of niche partitioning and evaluated the degree to which microbial taxa are specialized along the residence time gradient. Together, our findings contribute to a framework that aims to explain the maintenance of biodiversity in ecosystems subjected to the forces of physical turnover.

## METHODS

### Experimental design

We experimentally manipulated residence time across a set of chemostats (*n* = 49). The vessels had an operating volume of 40 mL (Fig. 1A). We used twice-autoclaved lake water as medium, which was collected from University Lake, a meso-eutrophic reservoir located in Griffy Woods, Bloomington, Indiana, USA (39.189° N, 86.503° W) (Wisnoski et al. 2020). We inoculated each chemostat with a microbial community derived from a mixture of lake water and sediment from University Lake. We then allowed the chemostats to equilibrate for 24 h before adjusting the flow rates using a combination of peristaltic pumps and manual pipetting across four experimental blocks. The chemostats were then incubated at 25 °C and homogenized with a magnetic stir bar. Because rates of resource input and immigration are often proportional to volume and flow rate, we added a suspension of lake sediment microbial community to each chemostat every day at a residence-time dependent concentration (1% of estimated total cell turnover per day). After 20 d, we destructively sampled the chemostats to measure microbial community structure and function.

**Fig. 1.**
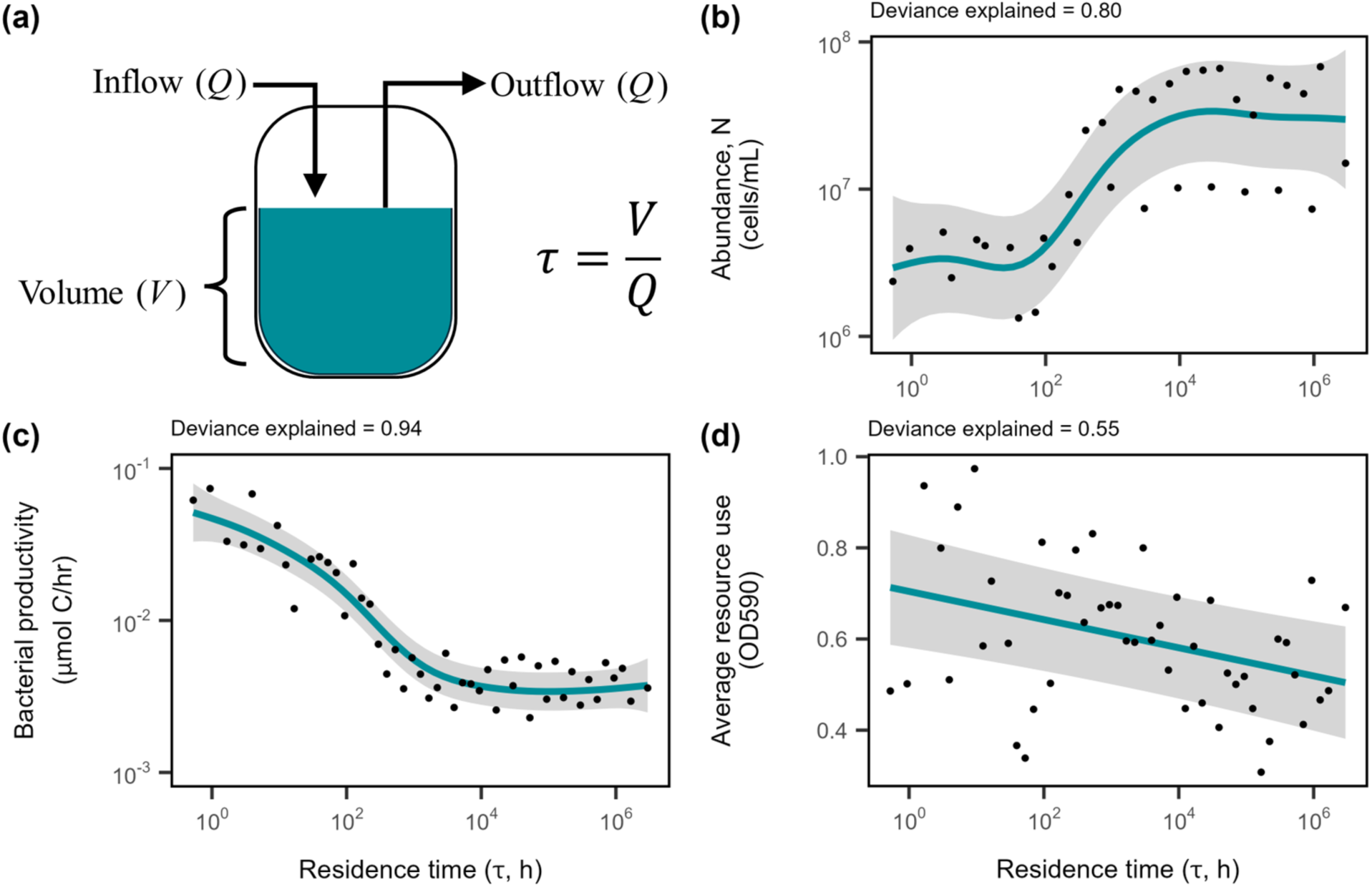
Abundance, productivity, and resource consumption of microbial communities along a residence time gradient. (A) We established a gradient of residence times by adjusting the flow rate (*Q*) of lake-water medium into well-mixed, fixed-volume (*V*) bioreactors (chemostats) that were inoculated with an aquatic microbial community. Generalized additive models describing the effect of residence time on **(B)** abundance (*N*), measured as cells/mL, **(C)** productivity, measured as the rate of biomass production (µmol C/hr) through a tritiated leucine assay, and **(D)** average resource consumption for each community measured via BioLog EcoPlates. Lines and shading represent fits and 95% CIs of GAM regressions, respectively. The percentage of deviance explained by the model is reported above each plot (Table 1).

**Table 1.** Effects of residence time on community structure. Response variables for each generalized additive model (Fit model) are shown with deviance explained and *P*-value for the total model (Model *P*-Value). Each smoothed term (Term) has an effective degree of freedom (edf) and *F*-value (*F*-statistic). Significance of each smoothed term (Term) was determined from term *P*-values (*P*-value) (α = 0.05). Restricted maximum likelihood values (REML) for each model are shown.

| Fit Model | Deviance explained | Model *P*-value | Term | Edf | *F*-statistic | *P*-value | REML |
| --- | --- | --- | --- | --- | --- | --- | --- |
| Abundance | 0.803 | 0.0001 | s($\tau$) | 4.69 | 17.04 | < 0.0001 | 13.2 |
|  |  |  | s(Set, bs = re) | 1.78 | 9.70 | 0.0002 |  |
| Bacterial Productivity | 0.940 | < 0.0001 | s($\tau$) | 5.04 | 96.97 | < 0.0001 | -22.4 |
|  |  |  | s(Set, bs = re) | 2.67 | 8.319 | < 0.0001 |  |
| Average Resource Use | 0.547 | 0.0003 | s($\tau$) | 1.00 | 13.00 | 0.0007 | -29.6 |
|  |  |  | s(Set, bs = re) | 2.76 | 12.69 | < 0.0001 |  |
| Resource PCoA Axis 1 | 0.40 | 0.0007 | s($\tau$) | 3.66 | 5.45 | 0.0007 | -37.5 |
| Resource PCoA Axis 2 | 0.68 | 0.0002 | s($\tau$) | 6.39 | 8.54 | < 0.0001 | -49.7 |
|  |  |  | s(Set, bs = re) | 2.17 | 2.84 | 0.0144 |  |
| Species Richness | 0.837 | 0.0093 | s($\tau$) | 5.06 | 18.23 | < 0.0001 | 257.7 |
|  |  |  | s(Set, bs = re) | 2.29 | 2.95 | 0.0186 |  |
| Species Evenness | 0.722 | < 0.0001 | s($\tau$) | 4.97 | 11.51 | < 0.0001 | -88.3 |
| Composition PCoA Axis 1 | 0.979 | 0.0055 | s($\tau$) | 7.88 | 119.01 | < 0.0001 | -38.7 |
|  |  |  | s(Set, bs = re) | 2.41 | 3.67 | 0.011 |  |
| Composition PCoA Axis 2 | 0.786 | < 0.0001 | s($\tau$) | 5.35 | 15.66 | < 0.0001 | -40.9 |
| Long $\tau$ Niche | 0.937 | < 0.0001 | s($\tau$) | 6.91 | 49.65 | < 0.0001 | -38.7 |
| Short $\tau$ Niche | 0.594 | 0.0007 | s($\tau$) | 5.68 | 5.22 | 0.0007 | -82.5 |

### Microbial Abundance

We measured microbial abundance using flow cytometry. After diluting samples 1:100 in phosphate buffered saline (PBS), we stained samples with 1 µL *Bac*Light™ RedoxSensor™ Green (RSG, Invitrogen, emission: 490/excitation: 520), a cell permeable electron transport chain activity indicator we used to gate live cells from cytometer noise. We left a replicate set of samples unstained, incubating all samples for 15 min in complete darkness at 37 °C. We fixed all samples with 25 µL of 25% glutaraldehyde and incubated them again for 30 min in complete darkness at 25 °C. The fixed cells were stored in the dark at 4 °C until they were run on a NovoCyte 3000 (Agilent, USA) with a 50 mW laser-emitting light at 488 nm. Green fluorescence from the RSG was measured on a logarithmic scale at 530/30 nm filter, set to a gain of 453 and a speed of 14 µL/min. We collected a total of 10,000 events with FSC-H larger than 300. We removed non-singleton events by gating on an SSC-H vs. SSC-A plot (Fig. S1A). Next, we gated for live cells on an RSG vs. count plot to remove cytometer noise seen in cell-free PBS samples while using the location of the unstained sample on the RSG axis to avoid removing live cells (Fig. S1B). We processed the data with NovoCyte software, NovoExpress (v. 1.4.1), using electronic gating, calculating absolute abundance as:

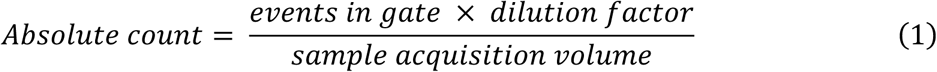

### Bacterial productivity

We measured the bacterial productivity (BP) in each chemostat using a tritiated (^3^H) leucine assay with replicate 1.5 mL samples (Smith and Azam 1992). We added ^3^H-leucine to a final concentration of 50 nM and then immediately stopped leucine incorporation in one replicate sample (kill-control) by adding 300 µL of 3 mM trichloroacetic acid (TCA). After 60 min of incubation for the remaining live samples (*n* = 3), we stopped the reactions with 300 µL of 3 mM TCA, removed unincorporated ^3^H-leucine, washed samples with 0.3 mM TCA, and added scintillation cocktail to all vials. The amount of radioactive leucine incorporated into bacterial proteins was measured with a Tri-Carb 2100TR Liquid Scintillation Counter (Packard Instrument Company). The measured counts per minute (CPMs) were converted to incorporation rates of µM C/h using estimates of cellular C per protein and the fraction of leucine in protein (Kirchman 2001).

### Resource consumption

We estimated changes in resource consumption for microbial communities along the residence time gradient with Biolog EcoPlates (Biolog, Hayward, CA, USA). Ecoplates contain 31 carbon sources along with a tetrazolium dye that changes color when reduced in the presence of NADH, which can be used to quantify the rate at which carbon substrates are consumed (Garland and Mills 1991). The 31-carbon sources can be grouped into seven higher-level categories: carboxylic acids, polymers, carbohydrates, amino acids, amines, esters, and phosphorylated carbon sources. We determined carbon consumption for each chemostat by measuring OD (590 nm) after a 48-h incubation at 25 °C and subtracting the water blanks.

Response values > 0.125 OD (590 nm) were counted as positive for consumption of a given carbon source (Garland 1997). We then calculated the mean rate of resource consumption across all carbon sources at each residence time. We used principal coordinates analyses (PCoA) to characterize the effects of residence time on resource consumption at 48-h. For our PCoAs, we used the Bray-Curtis dissimilarity metric from the ‘vegan’ package (version 2.6-4) in R (Dixon 2003).

### Microbial communities

#### Sequencing

We measured microbial diversity using high throughput 16S rRNA gene sequencing. We extracted genomic DNA from our Day 20 samples using the DNeasy UltraClean Microbial Kit (Qiagen, USA). After initial centrifugation, we incubated the bacterial resuspension in PowerBead Solution with lysozyme (50 µL 50 ng/mL) and 10 µL Proteinase K (20 mg/mL) for 1 h at 37 °C. DNA was eluted in 30 µL EB Buffer and stored at -20 °C. We amplified the V4 hypervariable region of the 16S rDNA gene using barcoded primers (515F and 806R) designed for the Illumina MiSeq platform (Caporaso et al. 2012). We used Phusion High Fidelity DNA Polymerase (New England BioLabs) with the 5X Phusion HF buffer, optional DMSO, and 2 µL template DNA. The conditions for the PCR reaction were 98 °C for 30 s, 30 cycles of 98 °C for 10 s, 68 °C for 30 s, and 72 °C for 15 s, followed by a final extension at 72°C for 10 min. We purified the sequence libraries using the AMPure Purification Kit (Bechman) and quantified the libraries with the Quant-it PicoGreen dsDNA kit (Invitrogen). We pooled the libraries to equal molar ratios (final concentration of 10 ng/library) and sequenced them using Illumina MiSeq 250x250 paired end reads (Reagents v2) at the Indiana University Center for Genomics and Bioinformatics.

#### Diversity and assembly

We processed raw 16S rDNA sequences to determine taxonomic composition of each chemostat along the residence time gradient. We assembled the paired-end raw 16S rDNA sequence reads into contigs, quality-trimmed, and aligned them to the Silva Database (version 138) (Quast et al. 2013). We detected and removed chimeric sequences using the VSEARCH algorithm (Rognes et al. 2016). We then split the sequences based on RDP taxonomy (Cole et al. 2009) then binned them into operational taxonomic units (OTUs) based on 97% sequence similarity. These initial sequence processing steps were completed using the software package mothur (version 1.48.0, Schloss et al., 2009). We also calculated phylogenetic distance between OTUs and generated a phylogenetic tree using the ‘dist.seqs’ method and Clearcut which implements a relaxed neighbor joining algorithm (Evans et al. 2006), respectively, both implemented in mothur. We removed OTUs with less than two reads across all samples. We quantified taxon richness (*S*) as the observed number of OTUs after rarefication and species evenness as Simpson’s evenness index (*D*^-1^/*S*) where *D*^-1^ is the inverse of Simpson’s diversity (Maurer and McGill 2011). We also characterized the effects of residence time on community composition with PCoAs, using the same methods as with resource consumption.

#### Niche partitioning

We tested whether there was a non-random degree of overlap in the relative abundance of OTUs along the residence time gradient using the Pianka index, a common measure of niche partitioning (Piana and Marsden 2012)

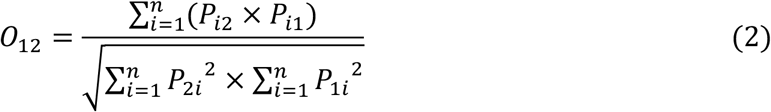

where *Pij* is the relative abundance of species *j* at residence time *i* and *n* is the total number of residence times (Pianka 1973). We first calculated the observed mean overlap in gradient use of OTUs found in >70% of sites (‘niche.overlap’ function in the ‘spaa’ R package, version 0.2.2) using method “pianka” (Pianka 1973, Zhang 2016). To test for significance, we then compared observed overlap in relative abundance of OTUs with values generated from null models. We generated two sets of 100 randomized chemostat x OTU matrices. In one set, all OTU abundances were drawn from a uniform distribution [0,1]. In the second set, observed zero abundances were preserved but all other elements of the matrix were drawn from a uniform distribution [0,1] (Winemiller and Pianka 1990, Gotelli and Graves 1996).

Next, we identified niches of OTUs that were non-randomly positioned along the residence time gradient. To accomplish this, we first calculated the Pearson’s correlation coefficient for each pair of OTUs. From those values, we then generated a distance matrix (where 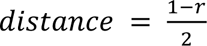) and performed *k*-means clustering by minimizing within-cluster variance (‘kmeans’ function from the ‘stats’ R package, version 4.3.2). To determine the appropriate number of niches, we plotted the within groups sum of squares by number of clusters. From this plot, we identified the number of niches where within group sums of squares was minimized without over-splitting (*n* = 5). We then calculated a combined relative abundance for each niche of OTUs at each residence time. To determine if the niches identified were phylogenetically clustered or randomly dispersed, we calculated the phylogenetic dispersion (*D*) of the two main niches using the ‘phylo.d’ function in the ‘caper’ package (version 1.0.3; Fritz and Purvis 2010) where *D* = 0 indicates phylogenetic clumping and *D* = 1 indicates random trait dispersion.

### Generalized additive models

We statistically modeled the effect of residence time on univariate microbial responses using generalized additive models (GAMs). GAMs are linear models where the response variable depends on smoothed functions of predictor variables, allowing for non-linear responses. We chose GAMs because, without having any *a priori* expectation of the predictive functions for our data, they accommodate flexible smoothing while still allowing interpretability as each predictive variable is encoded in the model. We used GAMs (‘gam’ function from the ‘mgcv’ R package, version 1.9-1) to fit bacterial abundance, BP, resource consumption, OTU richness, evenness, niche overlap, and relative abundance of niches, as a function of residence time using thin plate regression splines as the basis function (s(τ)) with a smoothing parameter (*λ*) estimated by restricted maximum likelihood (REML) (Wood 2017). We did not restrict the dimension of the basis used for the smoothing term (*k*). We report GAM *P*-values (α = 0.05) from a modified Wald test of whether the coefficient of the smoothed term is equal to zero (Wood 2013). We also report the deviance explained for each GAM. When experimental block was significant, we coded it with a random effect basis function (Experimental block, n = 4, s(Set, bs = “re”)). Models were selected based on a conditional Akaike information criterion (AIC) which applies a correction based on the effective degrees of freedom (edf) of the model. All statistical analyses were performed in R (version 4.3.2; R Core Team 2022).

## RESULTS

### Residence time altered microbial abundance and productivity

Bacterial abundance (*N*) increased nonlinearly with increasing residence time, plateauing at ∼3 x 10^7^ cells/mL (Table 1, Fig. 1B, *F4.69* = 17.0, *P* < 0.0001). In contrast, bacterial productivity (BP) declined non-linearly with increasing residence time, flattening out to a baseline of approximately 0.003 µmol C h^-1^ (Table 1, Fig. 1C, *F5.04* = 97.0, *P* < 0.0001).

### Residence time altered resource consumption

When averaged across all carbon substrates, resource consumption decreased near-linearly with increasing residence time (Fig. 1D, Table 1, *F1.00* = 13.0, *P* = 0.0007). When examined individually, we found that resource consumption significantly changed with residence time for 16 out of the 31 carbon substrates (Table S1, Fig. S5). In addition, residence time affected the composition of consumed resources. When visualized with a principal coordinates analysis (PCoA), the first two axes accounted for 81% of the total variation in consumed resources (Fig. 2A). The site scores from the first two axes of the PCoA both increased non-linearly with increasing residence time (Table 1, Fig. 2B, PCoA 1- *F3.66* = 5.4, *P* = 0.0007; Fig. 2C, PCoA 2- *F6.40* = 8.5, *P* < 0.0001).

**Fig. 2.**
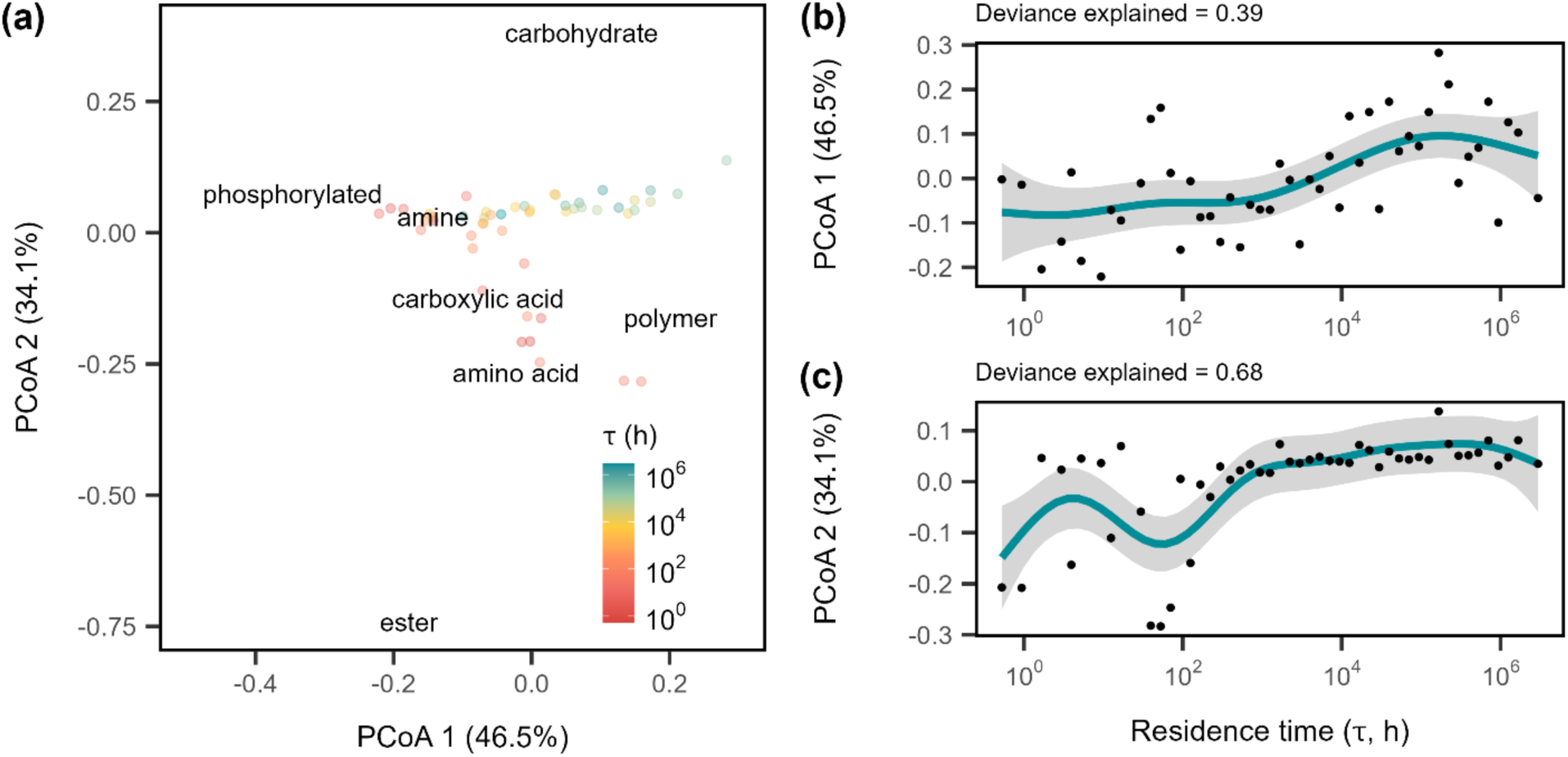
Resource use changed with residence time. (A) Principal coordinate analysis (PCoA) plot showing composition of resource use of 31 carbon sources of chemostat microbial communities at the end of 20 days of residence time treatment. Distances between circles represent dissimilarity between resource use profiles. Points are colored by τ. Average PCoA resource scores for each resource type are also shown on the plot. Generalized additive models show the effect of residence time on microbial community resource use along the first two axes of the principal coordinate analysis which account for **(B)** 47% and **(C)** 34% respectively. Lines and shading represent fits and 95% CIs of GAM regressions, respectively. The percentage of deviance explained by the model is reported above each plot (Table 1).

### Residence time altered microbial diversity

To characterize microbial diversity of the chemostats, we sequenced the 16S rDNA gene and recovered a total of 23,234 bacterial and 129 archaeal OTUs. Chemostats were dominated by seven phyla that are common to freshwater ecosystems: *Proteobacteria* (14-42%), *Planctomycetota* (11-45%), *Acidobacteriota* (4-24%), *Verrucomicrobiota* (5-14%), *Bacteroidota* (3-15%), *Actinobacteriota* (1-11%), and *Chloroflexi* (1-9%). Univariate measures of diversity were strongly affected by residence time. Taxon richness (*S*) increased non-linearly with increasing residence time (Fig. 3A, *F5.06* = 18.2, *P* < 0.0001) ranging from 2,000 to 6,000 OTUs, saturating at longer residence times. As residence time increased, Simpson’s evenness (*ED-1/S*) also increased and plateaued at an evenness of 0.08 (Fig. 3B, *F4.97* = 11.5, *P* < 0.0001). Similar patterns of community richness and evenness were found when data were analyzed as amplicon sequencing variants (ASVs) (Supplemental methods, Fig. S2, Table S1). Multivariate measures of diversity were also strongly affected by residence time. When visualized with principal coordinates analysis (PCoA), the first PCoA axis accounted for 42% of the variation in the dataset while the second axis accounted for 8% of the variation (Fig. 4A). Residence time had a strong effect on the compositional trajectory in ordination space (Fig. 4A). When chemostat PCoA scores along the first two axes were plotted against residence time, they increased non-linearly with increasing residence time (Table 1, Fig. 4B, PCoA 1, *F7.88* =119.0, *P* < 0.0001; Fig. 4C, PCoA 2, *F5.35* = 15.7, *P* < 0.0001).

**Fig. 3.**
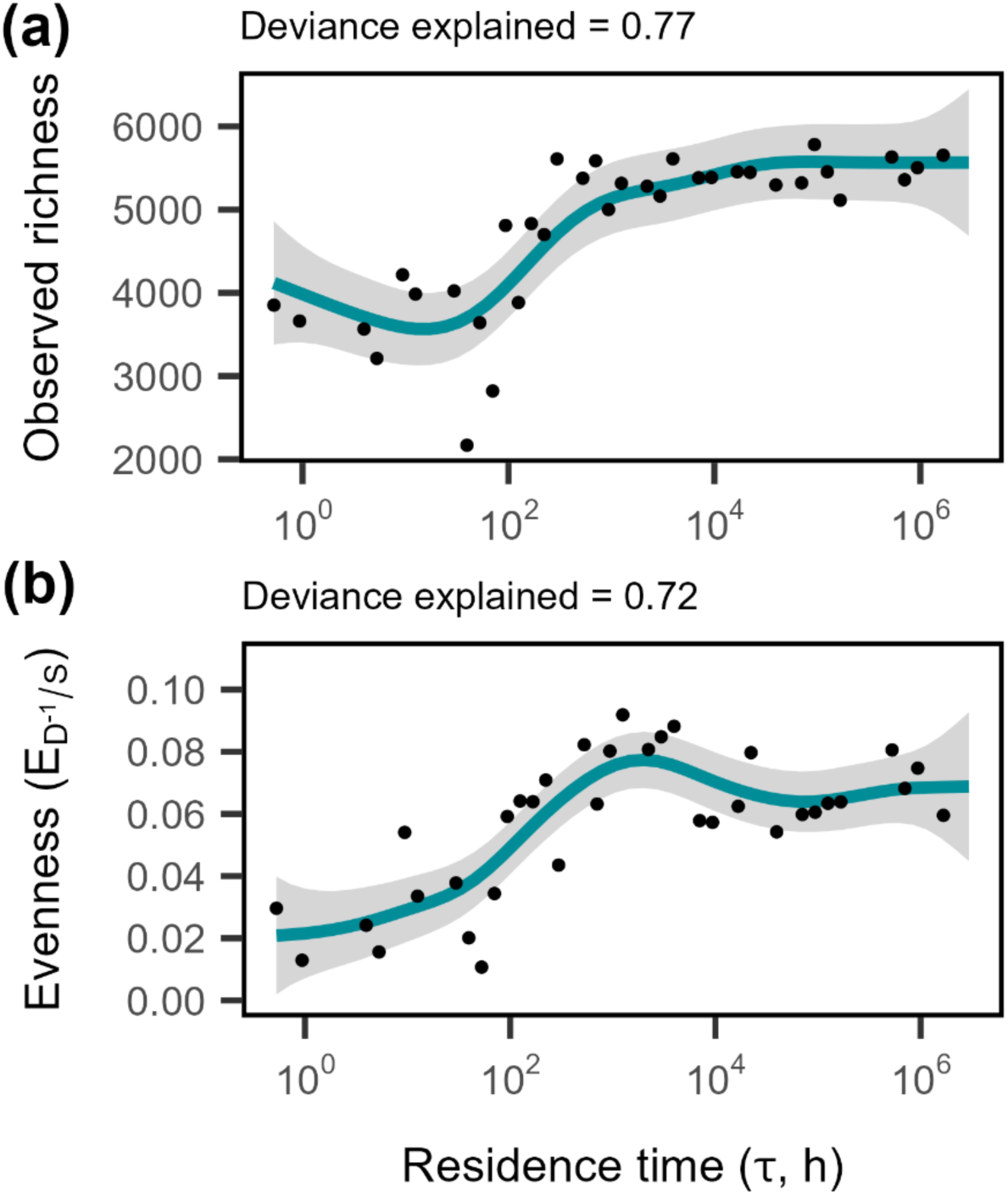
Microbial diversity increases with residence time. Generalized additive models describing the effect of residence time on **(A)** species richness (*S*), measured as observed number of OTUs, and **(B)** species evenness, calculated as Simpson’s measure of evenness (*E* = *D*^-1^/*S*; D^-1^ is the inverse of Simpson’s diversity). Lines and shading represent fits and 95% CIs of GAM regressions, respectively. The percentage of deviance explained by the model is reported above each plot (Table 1).

**Fig. 4.**
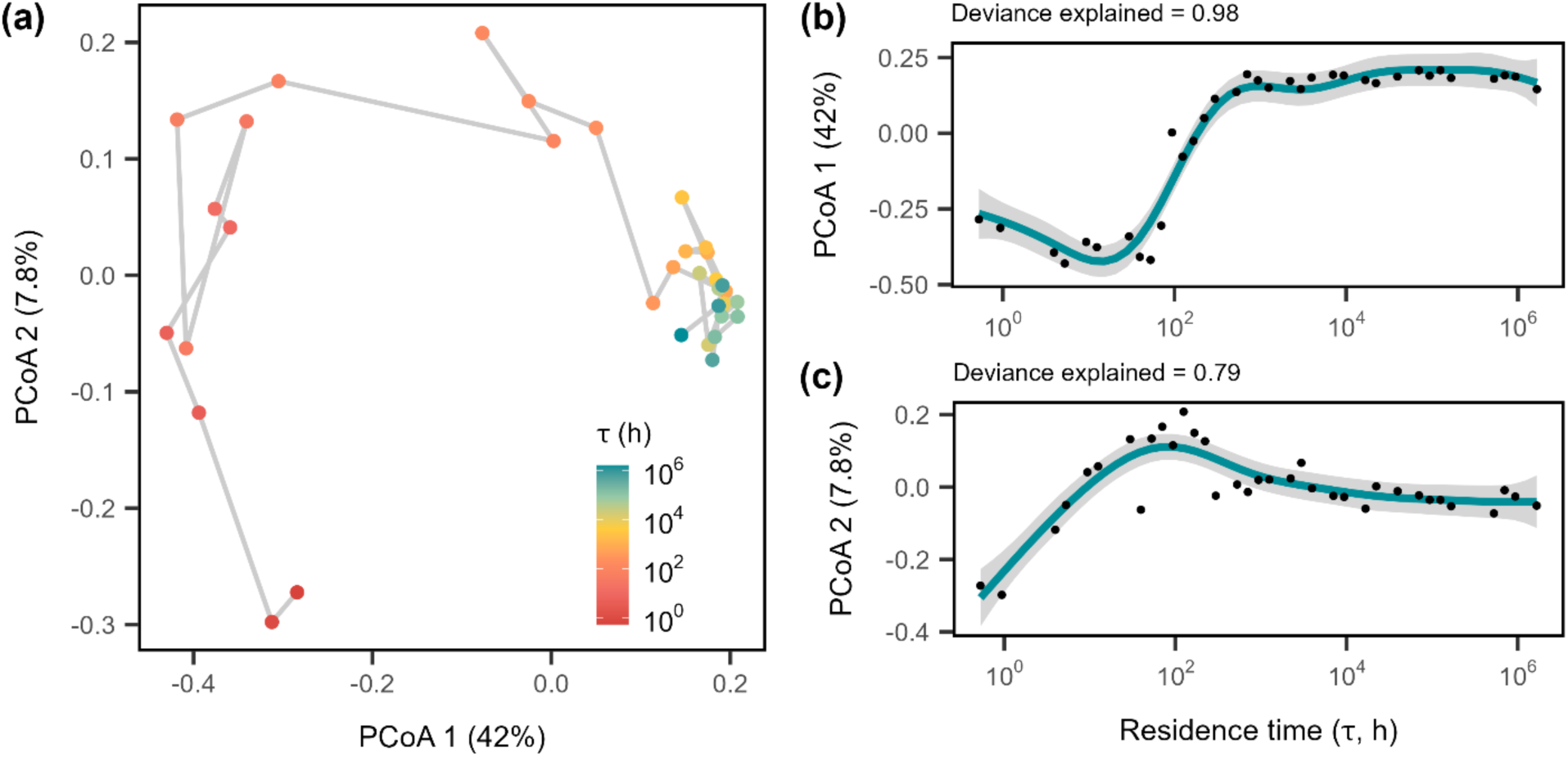
Community assembly along a residence time gradient. (A) Principal coordinate analysis (PCoA) plot showing how bacterial community composition responds to residence time after 20 d. Points are colored by τ and are connected by a line indicating the relationship between the residence times. Generalized additive models show the effect of residence time on microbial community composition along the first two axes of the principal coordinate analysis which account for **(B)** 42% and **(C)** 7.8% of total variation, respectively. Lines and shading represent fits and 95% CIs of GAM regressions, respectively. The percentage of deviance explained by the model is reported above each plot (Table 1).

### Residence time resulted in niche partitioning

Based on Pianka’s overlap index, there was significantly less overlap in microbial taxa along the residence time gradient than expected under null models (Fig. S3). Using *k*-means clustering and the minimization of within groups sum of squares, taxa fell into five niches (Fig. S4A & B, niche 1: *n* = 673, 30.5%; niche 2: *n* = 427, 19.4%; niche 3: *n* = 336, 15.3%; niche 4: *n* = 287, 13.0%; niche 5: *n* = 481, 21.8%). Two of these niches, niche 4 and niche 1, accounted for ∼45% of the total relative abundance and were positioned at short and long residence times, respectively (Fig. 5). Both niches exhibited phylogenetic clumping, but the short τ niche had a stronger phylogenetic signal (*D* = 0.38, *P*Brownian < 0.0001, *P*Random < 0.0001) than the long τ niche (*D* = 0.58, *P*Brownian < 0.0001, *P*Random < 0.0001). The two main niches were made up of 133 total orders, 14 unique at short τ, 75 unique at long τ, and 44 shared (Table S2).

**Fig. 5.**
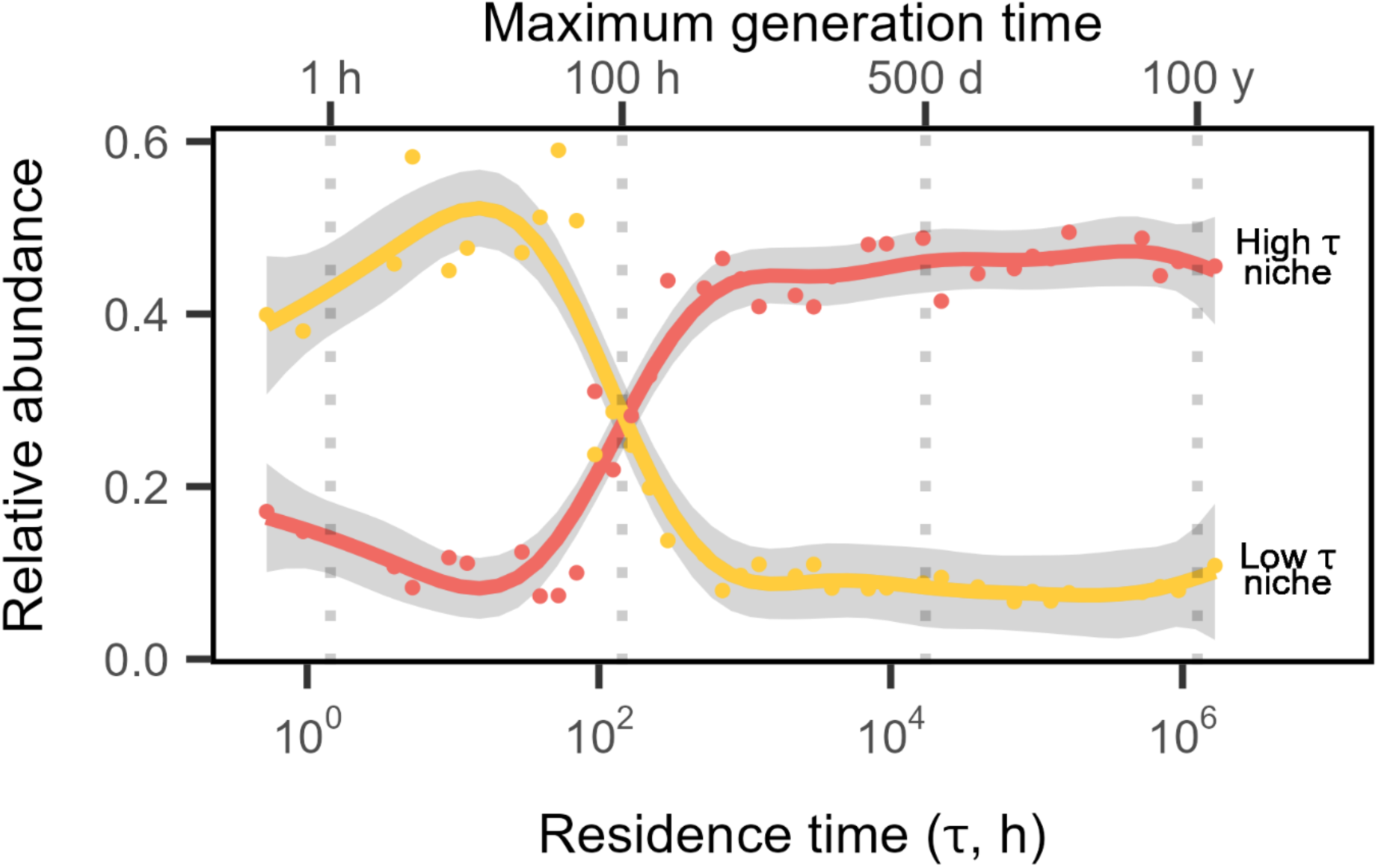
Niche partitioning along a residence time gradient. Total relative abundance of the two major niches identified through k-means clustering is shown along the residence time axis. The long τ niche, shown in red is identified as ‘niche 1’ in the text, and the short τ niche, shown in yellow, is identified as ‘niche 4’ in the text. Vertical grey dotted lines mark the estimated generation times (top axis) based on inverse of τ being equivalent to growth rate of taxa that are at equilibrium after 20 d. Lines and shading represent fits and 95% CIs of GAM regressions, respectively. The percentage of deviance explained by the model is reported above each plot (Table 1).

## DISCUSSION

We experimentally tested how residence time affects the assembly, diversity, and function of a complex microbial community using an array of continuous flow reactors (i.e., chemostats).

Patterns in bacterial abundance and productivity support the hypothesis that residence time structures communities through the opposing selective pressures of washout and resource limitation. These effects altered community assembly and resource consumption in ways that led to niche partitioning along the residence time gradient. The phylogenetic distribution of taxa within the two major niches suggests that conserved traits may contribute to patterns of diversity and function that emerge when communities are exposed to variation in physical turnover.

### Opposing selective forces: washout vs. resource limitation

Theory predicts that washout is the primary factor constraining abundance and species richness at short residence times (Locey and Lennon 2019). Under such conditions, individuals are removed more quickly than they can be replaced via reproduction. In support of this, we observed lower bacterial abundances and lower taxa richness in chemostats with short residence times (Fig. 1B & 3A). For the shortest residence time tested, a bacterium would have needed a growth rate of at least 1.9 h^-1^, which is equivalent to a doubling time of ∼20 min. There are many clinically relevant strains of bacteria that are capable of replicating on this timescale, including *Escherichia coli*, *Klebsiella pneumoniae*, and *Salmonella enterica* (Silva et al. 2009, Irwin et al. 2010, Liao et al. 2011). However, most environmental microorganisms do not divide this quickly. Based on genomically inferred doubling times, the vast majority of aquatic bacteria would not be able to persist in chemostats with short residence times (Weissman et al. 2021).

Therefore, while some microorganisms may avoid washout through fast growth rates, other bacteria may need to invest in traits such as adhesion and motility to persist in systems with short residence times.

In contrast, theory predicts that at longer residence times, resource limitation becomes the dominant ecological force, constraining metabolism, abundance, and species richness (Locey and Lennon 2019). In support of this, bacterial productivity and resource consumption declined in chemostats with increasingly long residence times (Fig. 1C & D). As a result, per capita rates of productivity were extremely low. Assuming an average cellular biomass of 50 fg (Cermak et al. 2017), we estimate that the doubling times of microorganisms in the long τ chemostats may be as high as 280 d. Under such conditions, bacteria must conserve internal resources to meet their maintenance energy requirements and prevent starvation (Jones and Lennon 2010). When cells die, their biomass components can be recycled, potentially supporting the metabolism of surviving individuals. For example, in a comparative analysis of phylogenetically diverse soil bacteria, nearly all populations could survive in the absence of exogenous resource inputs for at least 1,000 days (Shoemaker et al. 2021). The ability of microorganisms to persist under these conditions is critical given that most habitats on Earth have long residence times that lead to vanishingly low rates of resource supply (Basu and Pick 1997, Ambrosetti et al. 2003, Jönsson et al. 2004).

### Niche partitioning along the residence time gradient

Niche theory predicts that species must be sufficiently similar to coexist in a given environment, yet different enough to avoid competitive exclusion (Levine and HilleRisLambers 2009). When communities are structured to minimize niche overlap, clusters of taxa can emerge that specialize on different sets of conditions (Johnson et al. 2006, Dumbrell et al. 2010). This clustering reflects niche partitioning, a process that allows diverse groups of organisms to survive and coexist along environmental gradients. In the context of residence time, communities may be structured by multiple covarying variables including resources and physical factors like sheer stress generated by flow rate. Given its multifaceted impact, residence time may create an abundance of complex niches as environmental gradients overlap and interact.

In our experiment, we found evidence for niche partitioning. There was less overlap in the relative abundances of taxa across the residence time gradient than expected compared to null models (Fig. S3). Despite the wide range of residence times spanning many orders of magnitude, we identified only two dominant niches that contained half of all the individuals sequenced in our study. One niche comprised taxa specializing in short residence times, while the other comprised those specializing in long residence times. Some of the taxa found in the short τ niche belonged to the Pseudomonadota and Bacteroidota, which have been shown to increase at short τ (Vuono et al. 2015). Taxa belonging to Flavobacteriaceae were less abundant at short τ compared to long τ. However, the persistence of these taxa at short τ, despite the reduced number of species, suggests an ability to establish a population even under conditions that could otherwise lead to washout (Bulseco et al. 2024). In contrast, the long τ niche had a wider and asymmetric distribution. It was composed of more taxa than the short τ niche, but each had a lower relative abundance within the cluster (Fig. 5 & Table S2). The dominant taxa in this niche belonged to the Chloroflexota and Actinomycetota, consistent with observations made following a residence-time manipulation in a full-scale activated-sludge wastewater treatment plant (Vuono et al. 2015).

This distribution of taxa along the residence time gradient most likely reflects the underlying distribution of organismal traits (Locey and Lennon 2019). At short τ, fast growth rates may enable some taxa to maintain populations, but only a few traits, such as adhesion and motility, could reduce washout for slower-growing taxa (Ballyk et al. 1998). Alternatively, at long τ, there may be different ways for taxa to withstand resource limitation. For example, the number of enzyme classes present and the fractional share of rarer enzymes increased non-linearly with residence time in wastewater microbial communities (Mansfeldt et al. 2019). This may help explain the two-fold difference in number of taxa in the two niches (Table S2). We also see evidence for differential resource use for communities composed of taxa mainly found in the short τ vs. long τ niche (Fig. 2A & C). Furthermore, differences in trait distributions along the residence time gradient might be reflected by phylogenetic patterns, assuming that related taxa are more likely to share similar traits (Martiny et al. 2015). We found that taxa within a niche cluster were more phylogenetically related to one another than expected by chance. Additionally, there was a stronger phylogenetic signal for the taxa associated with the short τ niche than those associated with long τ.

To determine what traits are driving the niche partitioning, future studies should measure functional traits that are hypothesized to be important in systems with different residence times. This could be accomplished by isolating strains from different residence times and directly comparing functional traits such as growth rates, motility, resource use, and biofilm production (Lennon et al. 2012). However, cultivation bias toward fast-growing taxa might make this difficult at long τ. Alternatively, shotgun metagenomic sequencing could be used to perform a functional analysis, identifying relevant suites of genes and their relationship to residence time (Palanisamy et al. 2023).

### Applying residence time theory to natural systems

Residence time is associated with a wide range of biological phenomena in natural and managed ecosystems. In aquatic ecosystems, it is well known that rates of primary productivity and algal bloom emergence increase with residence time (Søballe and Kimmel 1987, Kim et al. 2022). Meanwhile, in host associated ecosystems, the severity of gastrointestinal disorders like Crohn’s disease increase with residence time and are associated with reduced gut microbial diversity (Ott et al. 2004, Fischer et al. 2017). However, additional factors must be considered when using residence time theory to understand dynamics in more complicated settings. While *V/Q* serves as a reasonable approximation of residence time in an idealized system, in reality there is a distribution of residence time. In a flowing system, some particles are removed faster than others due to chance and chaotic dynamics resulting from turbulence (Nauman 2008).

Variation can also arise when individuals evade washout by finding physical refugia or flow path sinks, allowing slow growing organisms to establish themselves in an otherwise fast-moving system (Drost et al. 2014). Residence time can also change over time, as seen in intermittent streams or changes in gut retention time in sick patients (Fischer et al. 2017, Blackman et al. 2021). Switching between states of washout stress and resource stress has the potential to prevent community responses to residence time from fully developing while also causing rapid changes in organismal abundance and species richness (Febria et al. 2012). Scaling up theory and findings from our small-scale reactors represents a future challenge for applying the residence-time framework to larger and more complex systems.

Despite these complications, residence time theory has potential for application in clinical, engineering, and environmental contexts. For example, in the gastrointestinal (GI) tract, residence time can be modified with water intake, laxatives, or dietary changes (Tropini et al. 2018). In cases where GI diseases select for harmful microorganisms and promote microbial overgrowth, manipulating residence time could provide an alternative intervention strategy that avoids the use of antibiotics (Procházková et al. 2023). In bioproduct manufacturing, residence time is critical for production rate and yield, but reactors typically operate with only one or two species (Fogler 2006). Understanding the effects of residence times on more diverse assemblages opens up opportunities for enhanced control over production processes, especially in light of recent insight into how stochastic and deterministic ecological forces influence the stability of synthetic microbial consortia (Li and Müller 2023). Last, eutrophication management is of concern to public health due to the increasing prevalence of illnesses associated with harmful algal blooms (HABs) in freshwater and coastal ecosystems (Grattan et al. 2016). Some HAB traits, such as toxin production and nitrogen fixation, become more prevalent with increasing residence time (Zhao et al. 2022). If flow rates (*Q*) can be adjusted based on an understanding of residence time’s influence on algal abundance and productivity, then it may be possible to mitigate algal blooms and their harmful effects (Cui et al. 2023). Future work should build on the whole-community approach used here to investigate the traits that drive niche partitioning, with the goal of developing a comprehensive understanding of how residence time affects communities in diverse ecosystems.

## ACKNOWLEDEMENTS

Research was supported by the National Science Foundation (DEB-1934554 to JTL; DBI- 2022049 to JTL), US Army Research Office Grant (W911NF-22-1-0014; W911NF-22-S-0008 JTL) and the National Aeronautics and Space Administration (80NSSC20K0618 to JTL). Code to reproduce all analyses is available on GitHub: https://github.com/LennonLab/residence-time-test.

