## Supplementary Information for "Residence time structures microbial communities through niche partitioning"

### SUPPLEMENTAL INFORMATION – METHODS

***Experimental design*** — For residence times < 41 h (1.7 d), we used peristaltic pumps to turnover the chemostats. For residence times > 41 h (1.7 d), we instead manually added and removed volume from the chemostats once per day. We ran the chemostats in four experimental blocks.

***ASV Diversity and assembly*** — To determine the robustness of alpha diversity patterns with residence time to clustering vs. denoising methods, we also used DADA2 (version 1.16.0, Callahan et al. 2016) to generate amplicon sequence variants (ASVs). We filtered and quality-trimmed the raw 16S rDNA sequence reads and then merged the paired reads. We then removed chimeras and assigned taxonomy to the sequence variants using the Silva Database (version 138). We calculated observed ASV richness ($S$) and evenness ($D^{-1}/S$) after rarefication ($N$ = 21,515).

### SUPPLEMENTAL INFORMATION – TABLES

| Fit Model | Deviance Explained | Model *P*-value | Term | Edf | *F*-statistic | *P*-value | REML |
|---|---|---|---|---|---|---|---|
| ASV richness | 0.74 | < 0.0001 | s(τ) | 5.52 | 10.96 | < 0.0001 | 235.2 |
| ASV evenness | 0.83 | 0.0053 | s(τ) | 5.55 | 13.59 | < 0.0001 | -41.3 |
| | | | s(Set, bs = re) | 2.39 | 3.56 | 0.0105 | |
| Niche 2 | 0.62 | < 0.0001 | s(τ) | 4.11 | 9.06 | < 0.0001 | -79.4 |
| Niche 3 | 0.63 | < 0.0001 | s(τ) | 3.35 | 11.80 | < 0.0001 | -59.9 |
| Niche 5 | 0.95 | < 0.0001 | s(τ) | 7.04 | 61.39 | < 0.0001 | -46.4 |
| D-Galactonic Acid γ-Lactone | 0.38 | < 0.0001 | s(τ) | 2.85 | 7.26 | 0.0003 | -1.5 |
| D-Galacturonic Acid | 0.12 | < 0.0001 | s(τ) | 1.00 | 6.29 | 0.0156 | 6.5 |
| D-Glucosaminic Acid | 0.28 | < 0.0001 | s(τ) | 1.55 | 9.87 | 0.0007 | -34.0 |
| D-Xylose | 0.40 | < 0.0001 | s(τ) | 2.02 | 11.57 | < 0.0001 | -38.6 |
| γ-Hydroxybutyric Acid | 0.20 | < 0.0001 | s(τ) | 1.00 | 11.48 | 0.0014 | 18.6 |
| Glucose-1-Phosphate | 0.32 | < 0.0001 | s(τ) | 2.92 | 4.95 | 0.0028 | -11.4 |
| Glycyl-L-Glutamic Acid | 0.21 | < 0.0001 | s(τ) | 2.48 | 3.42 | 0.0231 | -62.1 |
| L-Arginine | 0.31 | < 0.0001 | s(τ) | 1.47 | 10.35 | 0.0003 | 6.5 |
| L-Asparagine | 0.54 | < 0.0001 | s(τ) | 3.79 | 10.44 | < 0.0001 | 13.0 |
| L-Serine | 0.41 | < 0.0001 | s(τ) | 4.04 | 5.19 | 0.0008 | 7.2 |
| L-Threonine | 0.45 | < 0.0001 | s(τ) | 4.86 | 5.17 | 0.0004 | -24.8 |
| Putrescine | 0.37 | < 0.0001 | s(τ) | 1.00 | 27.16 | < 0.0001 | 0.7 |
| Pyruvic Acid Methyl Ester | 0.50 | < 0.0001 | s(τ) | 3.88 | 8.41 | < 0.0001 | -3.2 |
| Tween 40 | 0.77 | < 0.0001 | s(τ) | 4.86 | 23.46 | < 0.0001 | -26.5 |
| Tween 80 | 0.64 | < 0.0001 | s(τ) | 4.27 | 13.74 | < 0.0001 | -10.7 |

**Table S1. Output from generalized additive models for supplemental figures.** Response variables for each generalized additive model (Fit model) are shown with deviance explained and

*P*-value for the total model (Model *P*-Value). Each smoothed term (Term) has an effective degree of freedom (edf) and *F*-value (*F*-statistic). Significance of each smoothed term (Term) was determined from term *P*-values (*P*-value) ($\alpha = 0.05$). Restricted maximum likelihood values (REML) for each model are shown.

| Niche | Phylum | Order | # OTUs |
|---|---|---|---|
| Short τ (*n* = 287) | Bacteroidota | Flavobacteriales | 8 |
| | Pseudomonadota | Acetobacterales | 9 |
| Long τ (*n* = 673) | Actinomycetota | Gaiellales | 28 |
| | | IMCC26256 | 10 |
| | | MC-A2-108 | 9 |
| | Chloroflexota | Anaerolineales | 18 |
| | | JG30-KF-CM66 | 6 |
| | | KDA-96 | 21 |
| | Latescibacterota | Latescibacterota | 6 |
| | Methylomirabilota | Rokubacteriales | 8 |

**Table S2. Some taxonomic orders are specialists for short τ or long τ.** Some orders are found exclusively in either the short τ or long τ niche containing a notable number of representative OTUs (> 5). These orders are grouped by phylum within each niche and the number of OTUs in the taxonomic cluster is reported. The total number of OTUs in the niche are also reported (Short τ niche – *n* = 287; Long τ niche – *n* = 673

### SUPPLEMENTAL INFORMATION – FIGURE CAPTIONS

**Fig. S1. Flow cytometry gating for microbial abundance. (A)** Cells were gated on an SSC-H v. SSC-A plot for single-celled organisms to remove aggregates and a portion of baseline cytometer noise from the sample. **(B)** On an RSG count plot, cells were gated for RSG activity. This gate was positioned to remove machine noise captured with unstained PBS samples while avoiding removal of cells that were found in the unstained sample, meaning that live and active cells, regardless of activity level, were captured with this gate.

**Fig. S2. ASV richness and evenness reflect OTU richness and evenness. (A)** ASV richness, calculated as observed ASVs, increases along the residence time gradient, following the same pattern as OTU richness (Fig. 3A) **(B)** ASV evenness, calculated as Simpson's measure of evenness ($E = D^{-1}/S$; $D^{-1}$ is the inverse of Simpson's diversity), also increases with residence time in a similar manner to OTU evenness (Fig. 3B). Lines and shading represent fits and 95% CIs of GAM regressions and percentage of deviance explained by the model is reported above each plot (Table S1).

**Fig. S3. Observed niche overlap was lower than expected from null models.** The first null model (Null model #1) pulled all chemostat-by-OTU relative abundances from a uniform distribution [0,1]. This null model had a mean Pianka's niche overlap of 0.76 with a Cohen's D of 175.19. The second null model (Null model #2) maintained chemostat-by-OTU relative abundances that were zero, pulling the rest of the abundances from a uniform distribution [0,1]. This null model distribution had a mean Pianka's niche overlap of 0.64 with a Cohen's D of

45.44. Both null model distributions showed higher niche overlap than what was observed across all chemostats and OTUs, which is a Pianka's niche overlap of 0.61.

**Fig. S4. Five niches were identified across the residence time gradient using k-means clustering. (A)** Within group sum of squares of clusters fit with a LOESS regression indicated five niches as the optimal number of clusters. **(B)** Total relative abundance of all five niches identified through k-means clustering is shown along the residence time axis. Lines and shading represent fits and 95% CIs of GAM regressions (Table 1 & S1).

**Fig. S5. Use patterns of individual carbon sources vary, but mainly decrease with increased residence time.** Individual carbon source use was measured with a colorimetric use assay (BioLog EcoPlate) as color formation measured at OD (590 nm) at 48-h. Only resources with significant relationships with residence time after Benjamini-Hochberg multiple testing corrections are shown. Lines and shading represent fits and 95% CIs of GAM regressions (Table S1).

**Fig. S1.**

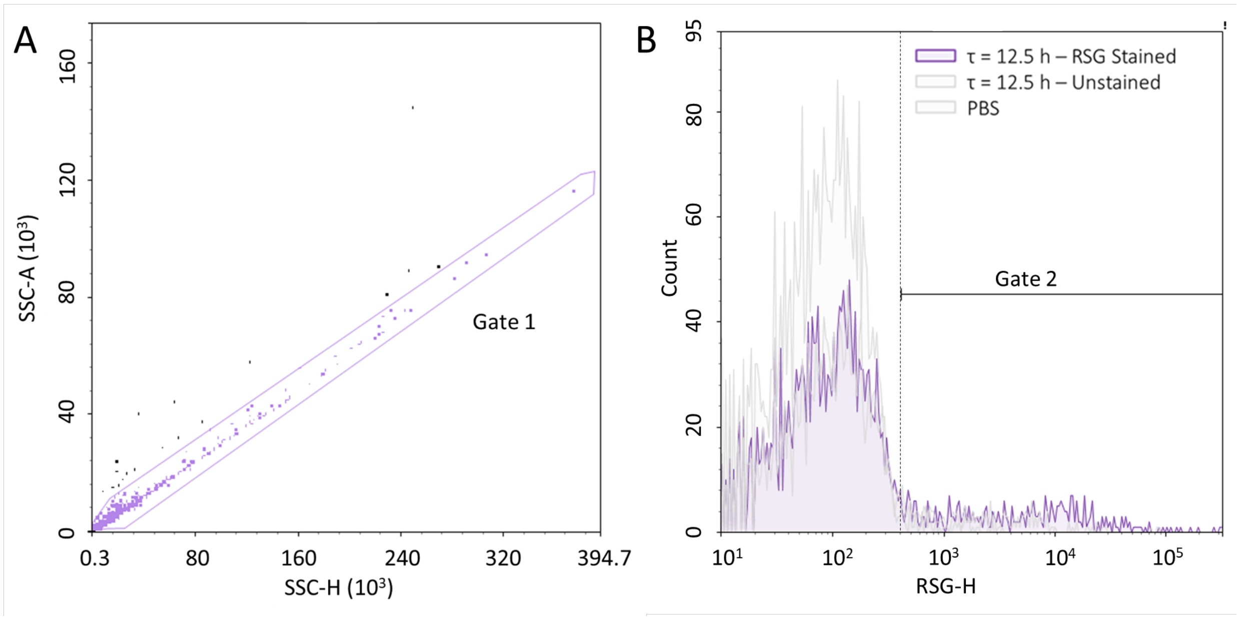
A
SSC-A (10³)
160
120
80
40
0
0.3
80
160
240
320
394.7
SSC-H (10³)
Gate 1
B
95
80
60
40
20
0
Count
τ = 12.5 h – RSG Stained
τ = 12.5 h – Unstained
PBS
Gate 2
10¹
10²
10³
10⁴
10⁵
RSG-H

**Fig. S2.**

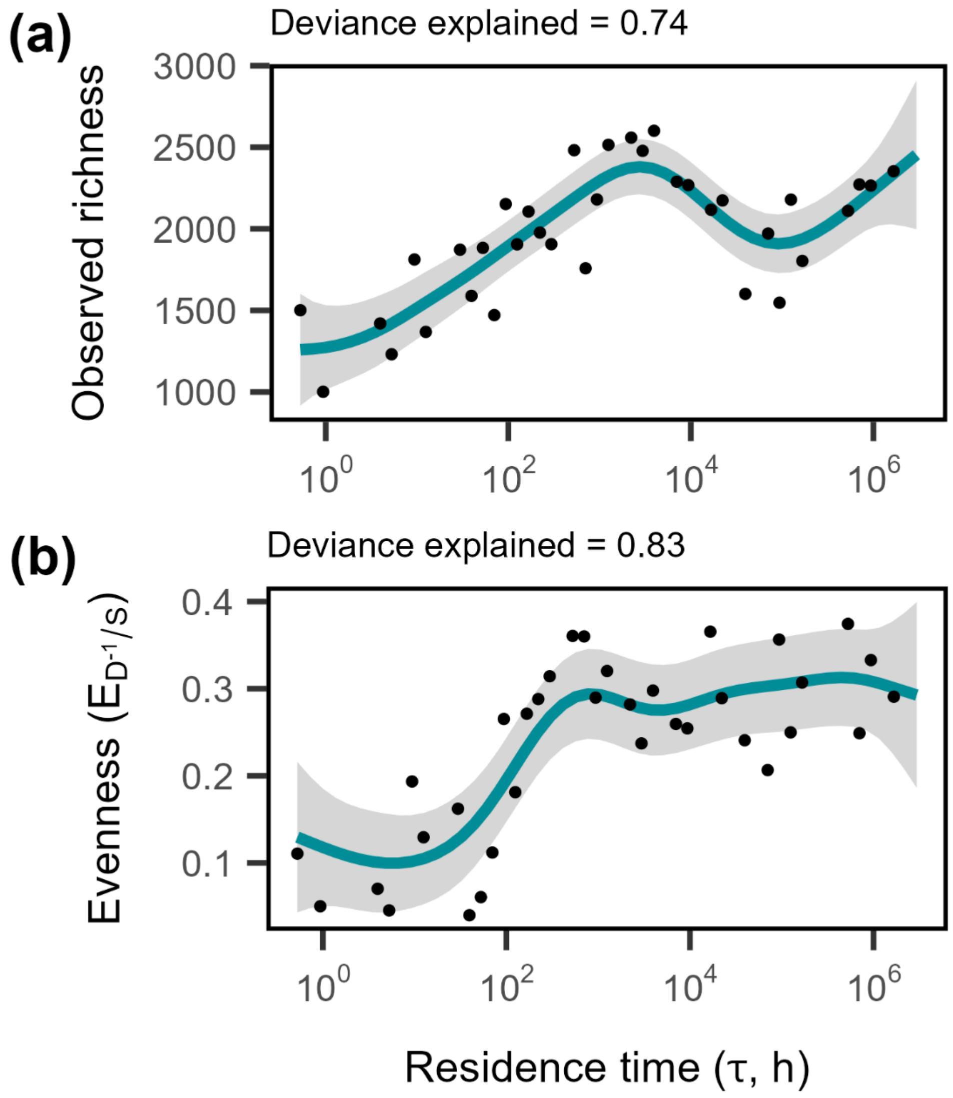
(a)
Deviance explained = 0.74
Observed richness
3000
2500
2000
1500
1000
$10^0$
$10^2$
$10^4$
$10^6$
(b)
Deviance explained = 0.83
Evenness ($E_{D^{-1}}/s$)
0.4
0.3
0.2
0.1
$10^0$
$10^2$
$10^4$
$10^6$
Residence time (τ, h)

**Fig. S3.**

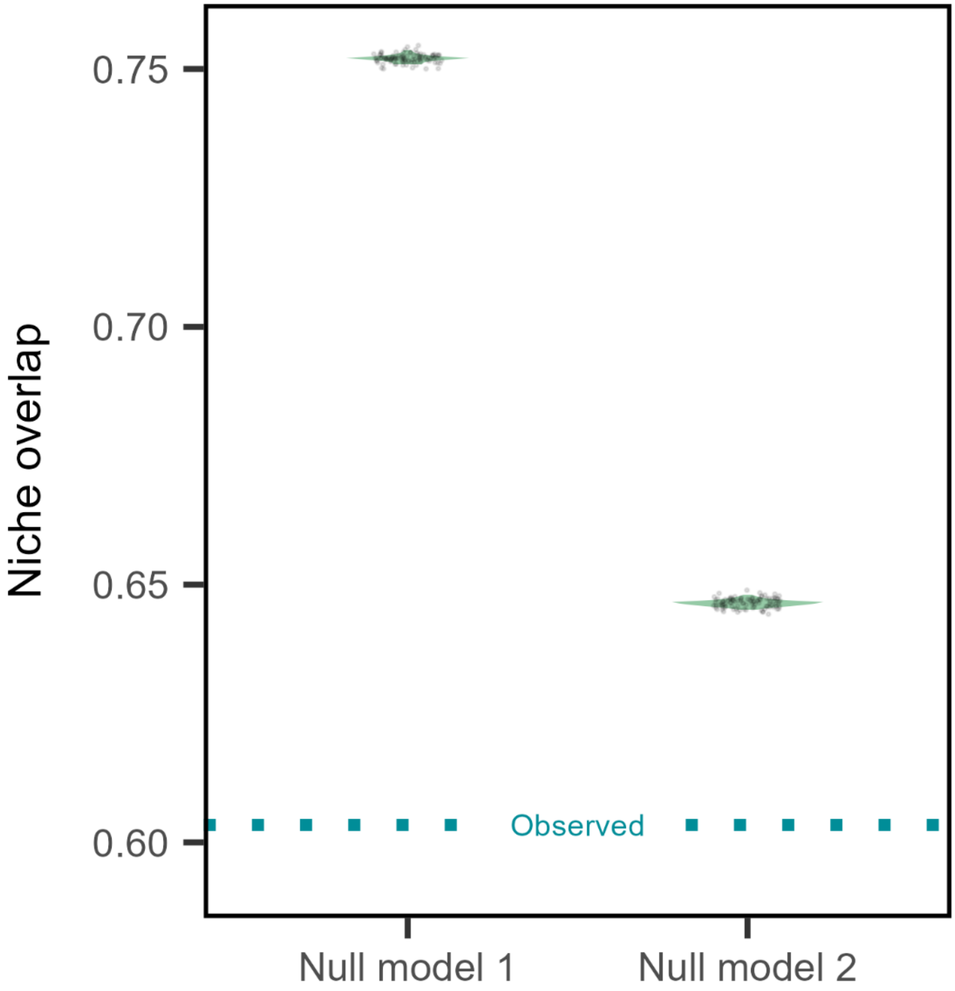

Niche overlap
0.75
0.70
0.65
0.60
Observed
Null model 1
Null model 2

**Fig. S4.**

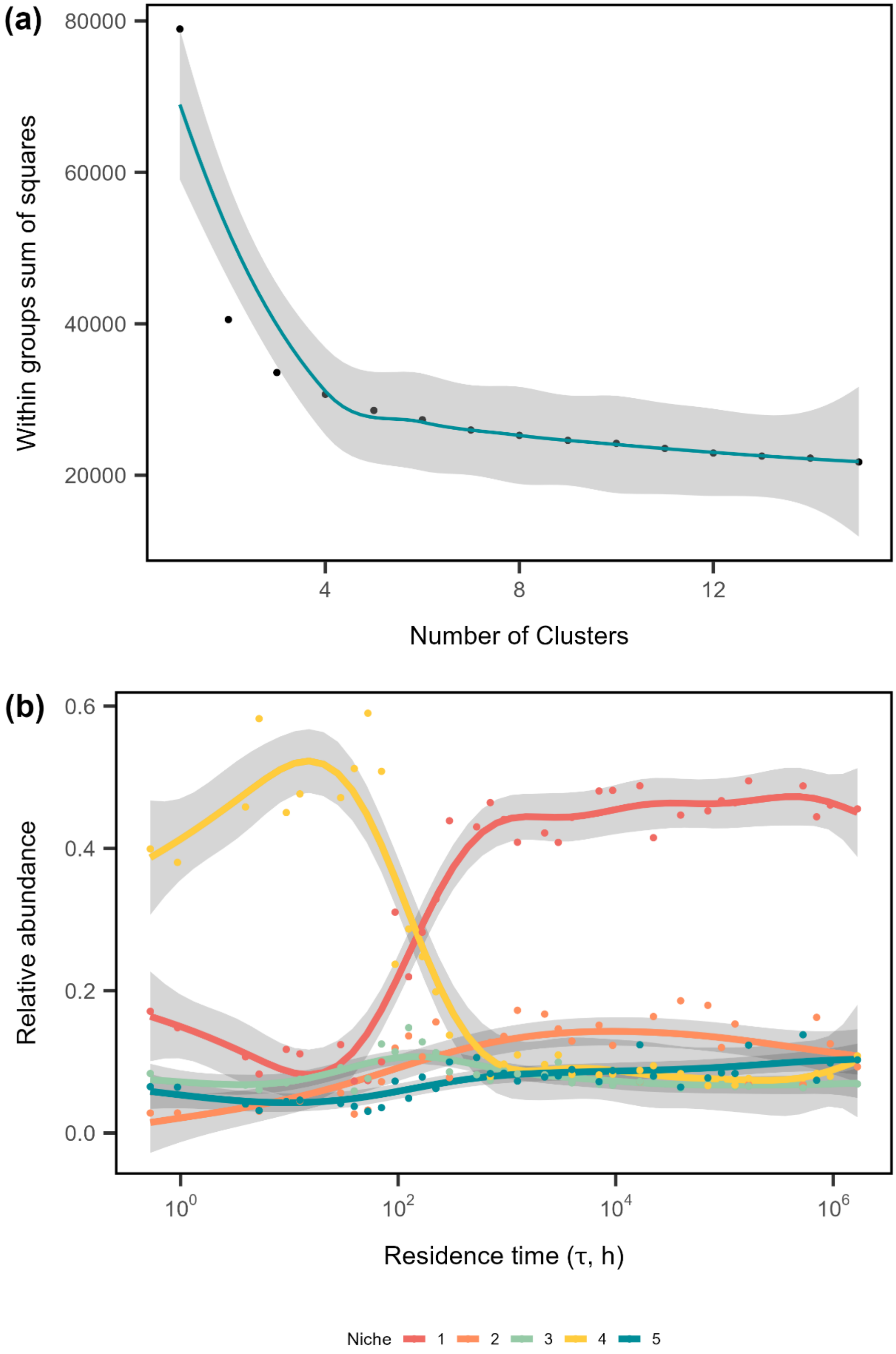
(a)
80000
60000
40000
20000
Within groups sum of squares
4
8
12
Number of Clusters
(b)
0.6
0.4
0.2
0.0
Relative abundance
$10^0$
$10^2$
$10^4$
$10^6$
Residence time (τ, h)
Niche 1 2 3 4 5

**Fig. S5.**

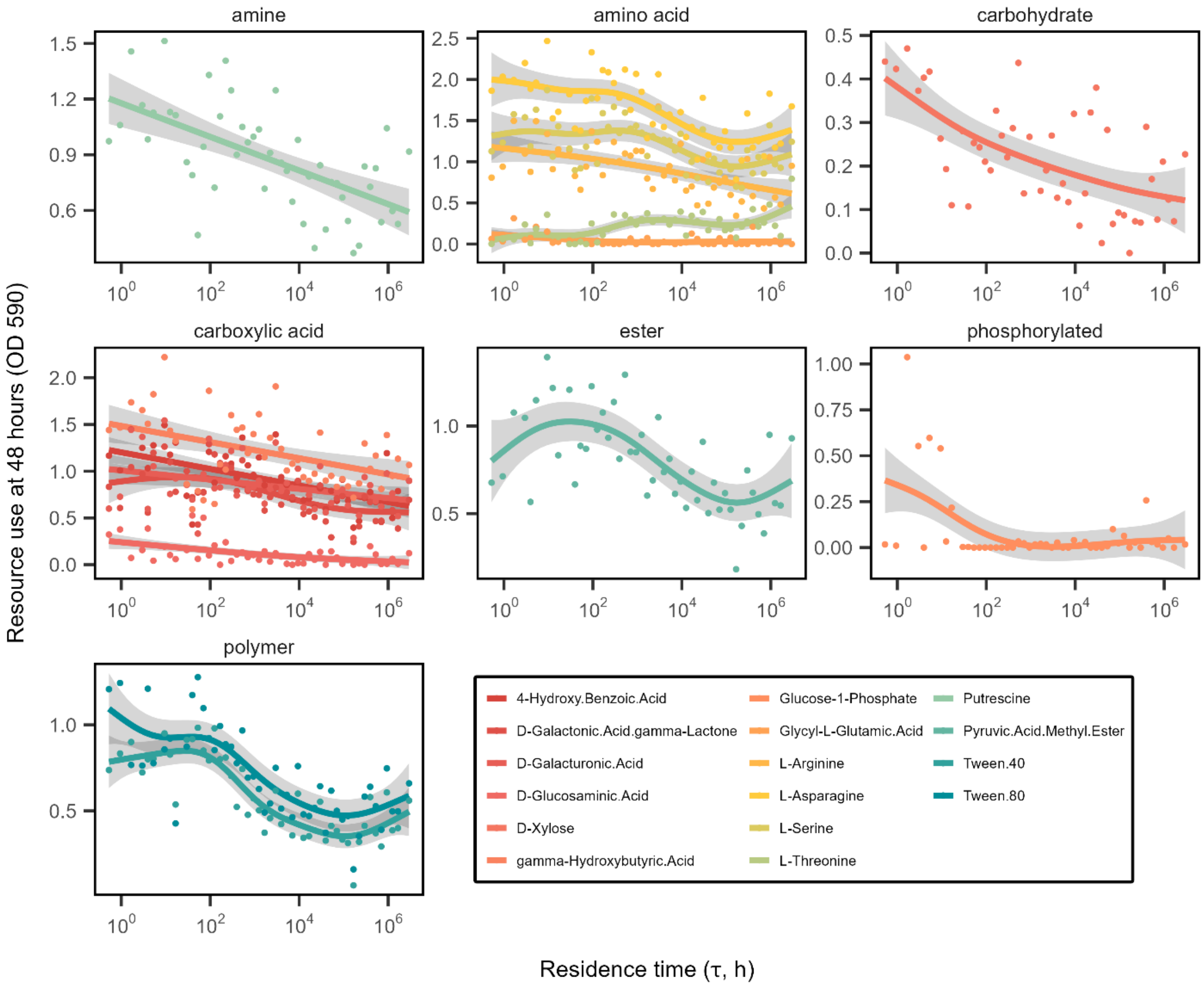
amine
amino acid
carbohydrate
carboxylic acid
ester
phosphorylated
polymer
Resource use at 48 hours (OD 590)
Residence time (τ, h)
4-Hydroxy.Benzoic.Acid
D-Galactonic.Acid.gamma-Lactone
D-Galacturonic.Acid
D-Glucosaminic.Acid
D-Xylose
gamma-Hydroxybutyric.Acid
Glucose-1-Phosphate
Glycyl-L-Glutamic.Acid
L-Arginine
L-Asparagine
L-Serine
L-Threonine
Putrescine
Pyruvic.Acid.Methyl.Ester
Tween.40
Tween.80
